# Cooperative activities of dICA69^N-BAR^ and dCIP4^F-BAR^ domain proteins regulate membrane tubule organization *in vivo*

**DOI:** 10.1101/2023.08.12.553109

**Authors:** Bhagaban Mallik, Sajad Bhat, Vimlesh Kumar

## Abstract

Intracellular membrane tubules play a crucial role in diverse cellular processes, and their regulation is facilitated by Bin-Amphiphysin-Rvs (BAR) domain-containing proteins. This study investigates the roles of dICA69^N-BAR^ and dCIP4^F-BAR^ *in vivo*, focusing on their impact on *in vivo* tubule organization. Through cell culture and immunofluorescence staining, we observed co-localization of endogenous dICA69 with dCIP4-induced membrane tubules, indicating their potential recruitment for tubule formation and maintenance. Additionally, dCIP4-positive tubules exhibit enrichment of actin regulatory proteins such as Wasp, SCAR, Arp2, Arp3, and Syndapin. Overexpressing dICA69^N-BAR^ in S2R+ cells reveals distinct punctate patterns in the perinuclear region. An earlier study indicated that F-BAR proteins spontaneously segregate from the N-BAR domain-containing proteins during membrane tubule formation. In contrast, our observation supports a model in which different BAR-domain family members can associate with the same tubule and cooperate to fine-tune the tubule width. Moreover, our analysis highlights how dCIP4^F-BAR^ facilitates the redistribution of dICA69^N-BAR^ punctae, leading to altered patterns within the cells. These cooperative activities of dICA69^N-BAR^ and dCIP4^F-BAR^ are vital for the precise organization of intracellular tubules. Understanding the underlying mechanisms governing this cooperation provides valuable insights into cellular dynamics and the organization of membrane tubules. The implications extend to various physiological and pathological conditions related to intracellular membrane dynamics.

## Description

BAR domain-containing proteins are crucial in membrane organization, dynamics, and tubule generation within the cells (Chernomordik and Kozlov, 2003; Kumar et al., 2009b; Mallik et al., 2022; Mallik et al., 2017; Taylor et al., 2019). Structural analysis reveals that the BAR domain possesses a characteristic crescent-like shape, enabling them to bind and induce curvature in cellular membranes (Boucrot et al., 2012; Gallop et al., 2006). BAR domain proteins actively participate in numerous cellular processes by sculpting the membrane, including endocytosis, membrane trafficking, and tubule formation (Baumgart et al., 2011; Wang et al., 2009). Through their ability to sense and shape membrane curvature, BAR domain proteins can facilitate the generation of tubular structures from flat membranes (Simunovic and Voth, 2015). These tubules are dynamic, essential for cellular compartmentalization, and serve as conduits for efficiently transporting various cargo, such as proteins and lipids.

The BAR domain proteins act as versatile membrane organizers and dynamic regulators, pivotal in maintaining cellular homeostasis and ensuring proper intracellular communication and function (Billcliff et al., 2016; Carman and Dominguez, 2018). Among various BAR domain-containing proteins, F-BAR (Formin-Binding Protein 1 Homology-Bin-Amphiphysin-Rvs) and N-BAR domains have emerged as potential membrane remodelers for inducing membrane dynamics and remodeling (Liu et al., 2015; Takano et al., 2008; Wang et al., 2009). F-BAR domain-containing proteins are typically involved in the early stages of membrane deformation, leading to the generation of long, tubular structures (Boucrot et al., 2012). However, N-BAR domains primarily function in the later stages of membrane remodeling, stabilizing the curved membrane and facilitating the formation of shorter, more rounded tubules (Shimada et al., 2007).

Despite several structural and biochemical studies, the *in vivo* interaction of N-BAR and F-BAR proteins remains elusive at the cellular level. Earlier cell culture study has identified dCIP4 as one of the potential molecules for tubule formation in cultured cells (Fricke et al., 2009). We endeavored to confirm the previously identified role of dCIP4^F-BAR^ and set out to characterize the interplay between dCIP4 (an F-BAR domain-containing protein) and dICA69 (an N-BAR domain-containing protein), essential for regulating the localization of dICA69^N-BAR^ perinuclear punctae and inducing membrane dynamics in S2R+ cultured cells. In order to understand the nature of dCIP4^FL^-induced membrane tubules, we immunostained transiently expressing S2R+ cells positive for dCIP4^FL^-EYFP with dICA69 antibodies. Our immunostaining result reveals that endogenous dICA69 is tightly located with the CIP4-positive membrane tubules (Figure 1, A-A′, B-B′). Consistent with the previous finding, overexpression of dICA69^N-BAR^ did not induce any detectable tubules. It formed punctuate-like structures at the perinuclear region of the cells (Figure 1, C-C′) (Mallik et al., 2017). Next, to understand the *in vivo* interaction of F-BAR and N-BAR domains, we co-express dICA69^N-BAR^ and dCIP4^F-BAR^ domains in S2R+ cells. Interestingly, the co-expression of the dCIP4^F-BAR^ domain results in the redistribution of dICA69^N-BAR^ punctae in the cells (Figure 1, D-D′). Thus, in contrast to the previous findings, our analysis revealed a novel observation elucidating the *in vivo* regulation of N-BAR (dICA69) and F-BAR (dCIP4) proteins that might contribute valuable insights governing the dynamics and organization of membrane tubules, which needs further investigation (Frost et al., 2008).

**Figure 1.**
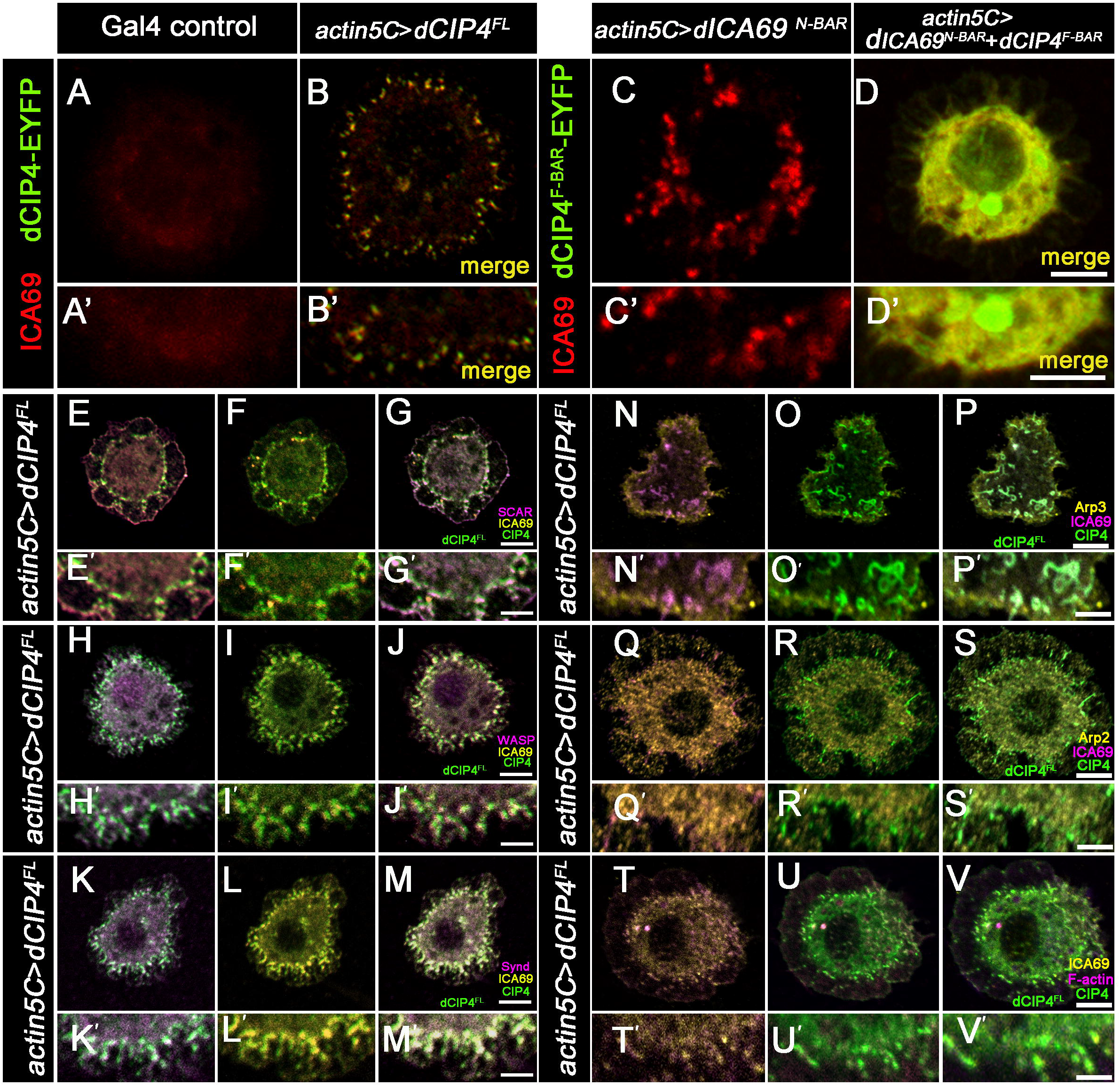
Overexpression of dCIP4^F-BAR^ redistributes dICA69 ^N-BAR^ punctae and is localized to endogenous dICA69 in S2R+ cells.: (A-D′) Confocal images of transfected *actin5C*-Gal4 control S2R+ cells incubated with transfection reagent (Mirus *TransIT*; A,A′), dCIP4^FL^ (B,B′), ICA69^N-BAR^ (C,C′) and ICA69^N-BAR^+CIP4^F-BAR^ (D,D′) co-labelled with ICA69 and anti-GFP antibodies. Scale bar: 10 µm (A-D, E-V); 4 µm (A′-V′). Note that overexpression of dCIP4^F-BAR^ leads to redistribution of dICA69^N-BAR^ perinuclear punctae, further resulting in altered patterns within the cells. Moreover, endogenous ICA69 is localized to CIP4-induced tubules. (E-M, E′-M′) Confocal images of cells transfected with full-length dCIP4^FL^ and immunolabelled with CIP4 (green), SCAR/Wasp/Synd (magenta), and ICA69 (yellow) antibodies. Note that SCAR, Wasp, and Syndapin are highly enriched at the CIP4-positive tubules. (N-Q, N′-Q′) Confocal images of cells transfected with full-length dCIP4^FL^ and immunolabelled with CIP4 (green), Arp3/Arp2 (yellow) and ICA69 (magenta) antibodies. Confocal images show that CIP4-positive tubules are highly enriched with Arp2/3 actin regulatory proteins. (T-V, T′-V′) Confocal images of cells transfected with full-length dCIP4^FL^ and immunolabelled with CIP4 (green), ICA69 (yellow) antibodies, and F-actin (magenta) dyes, respectively. Note that F-actin is highly enriched at the CIP4-positive tubules.

Further analysis revealed that dCIP4^FL^-induced membrane tubules show enrichment of Wasp, SCAR, Arp2, and Arp3, four of the positive regulators of actin polymerization (Figure 1, E, E′-J, J′, N, N′-S, S′). Additional immunostaining results showed that CIP4-positive tubules are tightly localized to Syndapin and F-actin (Figure 1, K, K′ - M, M′, T, T′-V, V′). These data suggest that dCIP4 relocalizes actin-regulatory proteins at the site tubule formation in S2R+ cells. Our results showed that dCIP4 and dICA9 proteins could be localized to the same membrane tubules, generating novel ideas for investigating N-BAR and F-BAR interactions *in vivo* (Figure 1). We further defined the role of the dCIP4^F-BAR^ domain in regulating the distribution of dICA69^N-BAR^ perinuclear punctae, suggesting the *in vivo* interaction of N-BAR and F-BAR proteins in the cells. Finally, we showed that CIP4-induced membrane tubules relocalize actin regulatory proteins, possibly to stabilize nascent membrane tubules (Figure 1).

More interestingly, N-BAR and F-BAR proteins play a crucial role in regulating actin dynamics at the plasma membrane through the interaction of actin-regulatory proteins such as Wasp (Wiskott-Aldrich Syndrome Protein) and NPFs (Nucleation promoting factors) (Fricke et al., 2009; Mallik and Kumar, 2017; Suetsugu and Gautreau, 2012; Takano et al., 2008). Recruitment of actin regulatory proteins induces actin polymerization, generating forces that help further deform the membrane and elongate tubules. Additionally, our previous report suggests that overexpression of dICA69 localizes actin regulatory proteins at the site of filopodia (Mallik et al., 2017). Consistent with this idea, our data suggest that endogenous dICA69 might relocalize Wasp, SCAR, at the site of dCIP4-induced membrane tubules, possibly through interaction with the actin regulators in S2R+ cells. Therefore, this actin-based pushing force, in coordination with the curvature-sensing ability of N-BAR and F-BAR proteins, enables efficient tubule extension and membrane remodeling during various cellular processes.

Additionally, earlier studies have indicated that N-BAR and F-BAR domain-containing proteins are segregated and localized to distinct tubules within the cells; more specifically, N-BAR proteins are localized to shorter, rounded, tighter tubules, whereas, F-BAR protein induces longer, elongated, and wider tubules (Boucrot et al., 2012; Frost et al., 2008). In contrast, we observed two distinct characteristics in the context of dCIP4^BAR^ and ICA69^N-BAR^ proteins; a) Overexpression of dCIP4^F-BAR^ redistributes dICA69^N-BAR^ perinuclear punctae resulting in an altered pattern within the cell, suggesting possible interaction of N-BAR and F-BAR proteins *in vivo*. b) dCIP4 and dICA69 are recruited to the same membrane tubules, further suggesting that a distinct set of N-BAR and F-BAR proteins can cooperate and form mixed protein scaffolds at the membrane surface, thus generating diversity in the width of membrane tubules. The formation of these heteromeric protein complexes possibly allows for enhanced curvature sensing and membrane tabulation.

Overall, the interplay between N-BAR and F-BAR proteins and their interaction with other cellular components form an intricate regulatory network that governs membrane dynamics, tubule formation, and actin regulation. These processes are essential for various cellular functions, including endocytosis, cell migration, and vesicular trafficking (Boulakirba et al., 2014; Hartig et al., 2009; Lemaigre et al., 2023). We do not know if dCIP4 and dICA69 proteins are involved in direct interaction within the cells or through indirect mechanisms to regulate tubule generation *in vivo*. This question may be of interest to future investigations. Further studies on the molecular mechanisms underlying N-BAR and F-BAR protein functions and their interactions will strengthen our understanding of fundamental cellular processes and may offer potential therapeutic targets for diseases associated with membrane trafficking and cell migration dysregulation.

## Material and Methods

### *Drosophila* S2R+ cell culture

*Drosophila* S2R+ cells were cultured in 1x Schneider’s *Drosophila* media (Invitrogen) supplemented with 10% fetal bovine serum (FBS), 50 U/ml penicillin, and 50 µg/ml streptomycin in 25-cm^2^ T-flasks (Corning) at room temperature. The cells (∼3x10^3^) were seeded in a six well-culture plate and transiently co-transfected with pUAST-ICA69^N-BAR^, pUAST-CIP4^N-BAR^, pUAST-CIP4^FL,^ and *actin5C*-Gal4 (1 µg each) using Mirus *TransIT* transfection reagent as described previously (Mallik et al., 2017; Mallik et al., 2023b).

### Antibodies and immunocytochemistry

For microscopic analysis, S2R+ cells co-expressing pUAST-ICA69^N-BAR^, pUAST-CIP4^N-BAR^, pUAST-CIP4^FL,^ and *actin5C*-Gal4 were adhered onto Concanavalin A (Sigma-Aldrich) coated coverslips and fixed with 4% formaldehyde solution for 15 minutes. Anti-ICA69 antibodies were used at 1:1000 dilutions (Mallik et al., 2017). The polyclonal antibodies anti-GFP (Roche), anti-Arp2, and anti-Arp3 were used at 1:200 dilutions. Other antibodies and dyes used were anti-Syndapin (Kumar et al., 2009a; Kumar et al., 2009b), anti-SCAR (Zallen et al., 2002), anti-Wasp (Bogdan et al., 2005), and Rhodamine conjugated phalloidin (Thermo Fisher Scientific) (1:200 dilutions). Anti-mouse and anti-rabbit Alexa fluor 488/568 conjugated secondary antibodies were incubated at 1:800 dilutions (Thermo Fisher Scientific). The cells were mounted in Vectashield (Vector Laboratory) and imaged with 63x/1.4NA objective in Zeiss LSM780 confocal microscope as described previously (Mallik et al., 2023a; Mallik and Frank, 2022; Raut et al., 2017; Sekhar et al., 2019).

## Authors contributions

<u>Bhagaban Mallik</u>, Conceptualization, Formal analysis, Investigation, Methodology, Project administration, Visualization, Writing-Original draft, Writing-Review, and editing^1*^. <u>Sajad Bhat,</u> Investigation, Writing-Review, and editing^1^.<u>Vimlesh Kumar</u>, Conceptualization, Formal analysis, Funding acquisition, Investigation, Methodology, Project administration, Resources, Visualization, Writing-Review, and editing^1*^.

## Acknowledgments

We thank Dr. Saumitra Dey Choudhury for his suggestions throughout this work and Dr. Nissi Varghese for helping with the initial experiments.

